# iPSC-derived neurons as a tool for probing molecular pharmacology of antipsychotic action

**DOI:** 10.1101/308486

**Authors:** Esther S. Kim, E. David Leonardo, Alex Dranovsky

**Affiliations:** Department of Psychiatry at Columbia University and the New York State Psychiatric Institute.

**Author notes:** Corresponding author contact information: Alex Dranovsky or David Leonardo, 1051 Riverside Dr. Unit 87, (AD) or 7105 (EDL), or.

## Abstract

**Background:** Induced pluripotent stem cell derived neurons (iPSC-Neurons) provide a potential way to investigate molecular mechanisms of psychotropic drug action in human neurons. Until now such studies have relied on animal models or artificial expression systems in transfected cells.

**Methods:** Induced pluripotent stem cells were subjected to a dual SMAD inhibition differentiation protocol. Resulting neurons were examined using qPCR, immunocytochemistry, viral transduction, and calcium imaging.

**Results:** Here we report the presence of target receptors for antipsychotic drugs in human iPSC-neurons. A cortical neuronal differentiation protocol resulted in cells that expressed D2, 5HT2A, and other target receptors. Moreover, stimulation with glutamate, dopamine, or the 5HT2A agonist DOI evoked calcium transients. We analyzed single cell responses, and found cells with signature response profiles to these ligands. In addition, pre-incubation of iPSC-neurons with clozapine altered the proportion of cells that responded to glutamate or DOI in a subpopulation of neurons.

**Conclusions:** Our results support the use of iPSC-neuron single cell pharmacology for studying how psychotropic medications modulate neuronal responses. Because these cells can be derived directly from patients, results derived from using iPSC-neurons have immediate relevance for personalized medicine.

**Significance Statement:** The current study examines the feasibility of using induced pluripotent stem cells from patients to generate neurons and study psychopharmacology. This article is broadly intended to inform the readership on the key points of iPSC-derived neurons as a system and how it can be used to understand antipsychotic pharmacology for potential clinical application. The specific advances include 1) demonstrating the presence of receptors targeted by antipsychotics on iPSC-derived neurons; 2) Using single cell analysis to identify human neurons with distinct responses to receptor modulation; and 3) Demonstrating that clozapine modulates glutamatergic and serotonergic responses in distinct human neuronal populations.

## Introduction

Since the seminal generation of induced pluripotent stem cells (iPSCs) from human somatic cells (Takahashi et al., 2007), numerous studies have examined iPSC-neurons from patients, including those with schizophrenia (Brennand et al., 2012), autism (Shcheglovitov et al., 2013; Liu and Scott, 2014), ALS (Kondo et al., 2014), Alzheimer’s (Mohamet et al., 2014), and Parkinson’s disease (Ryan et al., 2013). These cells express components of a fully functional neuron, including cytoskeletal and synaptic proteins (Shi et al., 2012; Espuny-Camacho et al., 2013), the ability to generate action potentials (Boulting et al., 2011), and calcium transients (Boulting et al., 2011; Dage et al., 2014). Thus far, studies have used iPSC-neurons to investigate changes in morphology, protein expression, and developmental trajectories in diseased states compared to controls. Another especially useful yet less explored platform to use this model is for studying mechanisms of psychotropic drug action.

Methods to understand how antipsychotics exert their effects have included human imaging studies, *in vitro* expression systems, and animal models. Positron emission tomography and single-photon emission computed tomography studies reveal antipsychotic binding sites and their binding capacities to dopamine and serotonin receptors in schizophrenia patients and other groups. These imaging studies are limited by the available radioligands and probe a limited number of receptors at a given time (Farde et al., 1992; Talbot and Laruelle, 2002). Moreover, thespatial and temporal resolutions of these techniques are not sufficient to identify cellular targets or reveal physiological mechanisms of action for these drugs at their receptors. Thus, to better understand cellular and molecular mechanisms underlying psychotropic action, stable cells lines, such as Chinese hamster ovary (CHO) cells and human embryonic kidney (HEK) cells with exogenous expression of receptors are often employed (Ruch et al., 1976; Schotte et al., 1996; Kuroki et al., 1999; Lawler et al., 1999; Burris et al., 2002; Kontkanen et al., 2002; Fell et al., 2012). However, given the highly-specialized organization of neuronal cell biology, the relevance of these *in vitro* systems to human psychopharmacology is limited. Because iPSCs can be generated from patients with known drug responses, they represent a potentially unique opportunity to study pharmacological mechanisms in an in vitro system with fewer hurdles to clinical translation. Furthermore, because iPSCs are stem cells, they have the capacity for extended self-renewal thereby providing an unlimited resource for mechanistic cell biology studies from each patient.

As stem cells, iPSCs can be differentiated into any cell of interest, including neurons, which is of particular importance in neurology and psychiatry, where access to live neurons from human subjects is limited. Many differentiation methods, including the current study, employ a monolayer cell culture, which allows for observation of single cell responses with the caveat that monolayers do not reflect the density and environment of neurons in the developing human brain. Methods for 3D culture have also been used, however, these methods provide limited access to individual cells and have other limitations. One unique strength of monolayer culture lies in providing the ability to observe live cellular and functional changes at single cell resolution for all cells within the culture. Some recent studies have examined mRNA expression and function of voltage gated calcium channels, GABA and glutamate receptors using such monolayer approaches (Brennand et al., 2011; Dage et al., 2014). These studies provided further support that iPSC-neurons exhibit characteristic physiological neuronal properties. However, the expression and functionality of receptors for modulatory amines like dopamine and serotonin has not been established in iPSC-neurons. By analyzing neuronal responses to psychoactive drugs in monolayer culture, we can examine cellular and molecular mechanisms by which antipsychotics exert their effects in iPSC-neurons. This method could potentially be used with patient iPSC-neurons to understand mechanisms underlying clinical drug responses.

Current antipsychotics all target dopamine D2 receptors (D2R), and this binding is correlated to clinical improvement in patients with schizophrenia (Nordstrom et al., 1993; Kapur et al., 2000). Antipsychotics also bind many additional receptors, for example the serotonin 5HT2A receptor (5HT2AR), raising speculations about other mechanisms of their clinical efficacy. Moreover, the diverse binding targets of antipsychotics can activate or block multiple intracellular signaling pathways, but the contribution of each receptor to drug action is unclear. Together the diversity of receptor and signaling targets for antipsychotics constitutes an exciting opportunity to develop new treatments and to individualize treatment for the currently available agents.

While in vitro systems are ideal for unraveling pharmacological differences between psychotropic drugs, they fall short on recapitulating the complexity of human neuronal receptor expression. However, with iPSC-neurons, we have the potential to probe antipsychotic effects on human neurons directly. In this proof of concept study, we investigated the cellular effects of antipsychotics on live human iPSC-neurons by monitoring their calcium responses. Changes in calcium levels can be used as a proxy of neuronal action potential activity (Kerr et al., 2005; Sasaki et al., 2008) and can thus serve as a first pass reporter for the cumulative cellular effects of receptor activation by medications. Moreover, live calcium imaging allows the monitoring of real time responses of many individual neurons at once with relative ease compared to, for example, electrophysiology. Using this system, we examined whether human iPSC-neurons express receptors that are targets of atypical antipsychotics and whether these receptors exhibit functional responses to neurotransmitters. Lastly, we examined whether the atypical antipsychotic clozapine, which has a unique clinical action profile compared to all other antipsychotics can modulate iPSC-neuronal responses to these neurotransmitters. Our results provide proof of principle that the iPSC-neuron system is a useful platform to study the pharmacology of antipsychotic action.

## RESULTS

iPSCs from a healthy subject were provided by the NIH CRM, and neurons were differentiated using the established Dual-SMAD inhibition protocol (Tomishima, 2008). This protocol generates neurons that resemble those of the forebrain, with the majority of the population consisting of glutamatergic neurons and smaller proportions of GABAergic and dopaminergic neurons (Chambers et al., 2009; Brennand et al., 2011). This mixed population of neurons was chosen because each subtype has been implicated in schizophrenia pathophysiology. iPSC-neurons were positive for neuronal markers MAP2 and Tuj1 via immunostaining (Fig 1A-D), and also positive for human Synapsin using a lentivirus reporter (Suppl Fig 1). We used CamKII immunoreactivity to identify glutamatergic principle neurons and found that the majority of iPSC-neurons (~80%) were CamKII positive (Fig 2A-E). This was further supported by expression of Vglut2 mRNA in the harvested cells (Suppl Fig 2). As described by others, using this differentiation protocol, a smaller proportion of GABAergic and dopaminergic neurons were also present, as identified by somatostatin (SS) and tyrosine hydroxylase (TH) immunostaining (Fig 2). Despite this, we did not detect significant levels of TH, GAD1, and GAD2 mRNA (Suppl Fig 2), likely due to the low proportion of these cells in our culture. Together these results indicate that as reported by others, Dual-SMAD inhibition results in differentiation of iPSCs into a heterogeneous population of predominantly principle glutamatergic with few dopaminergic and inhibitory GABAergic iPSC-Neurons.

**Figure 1.**
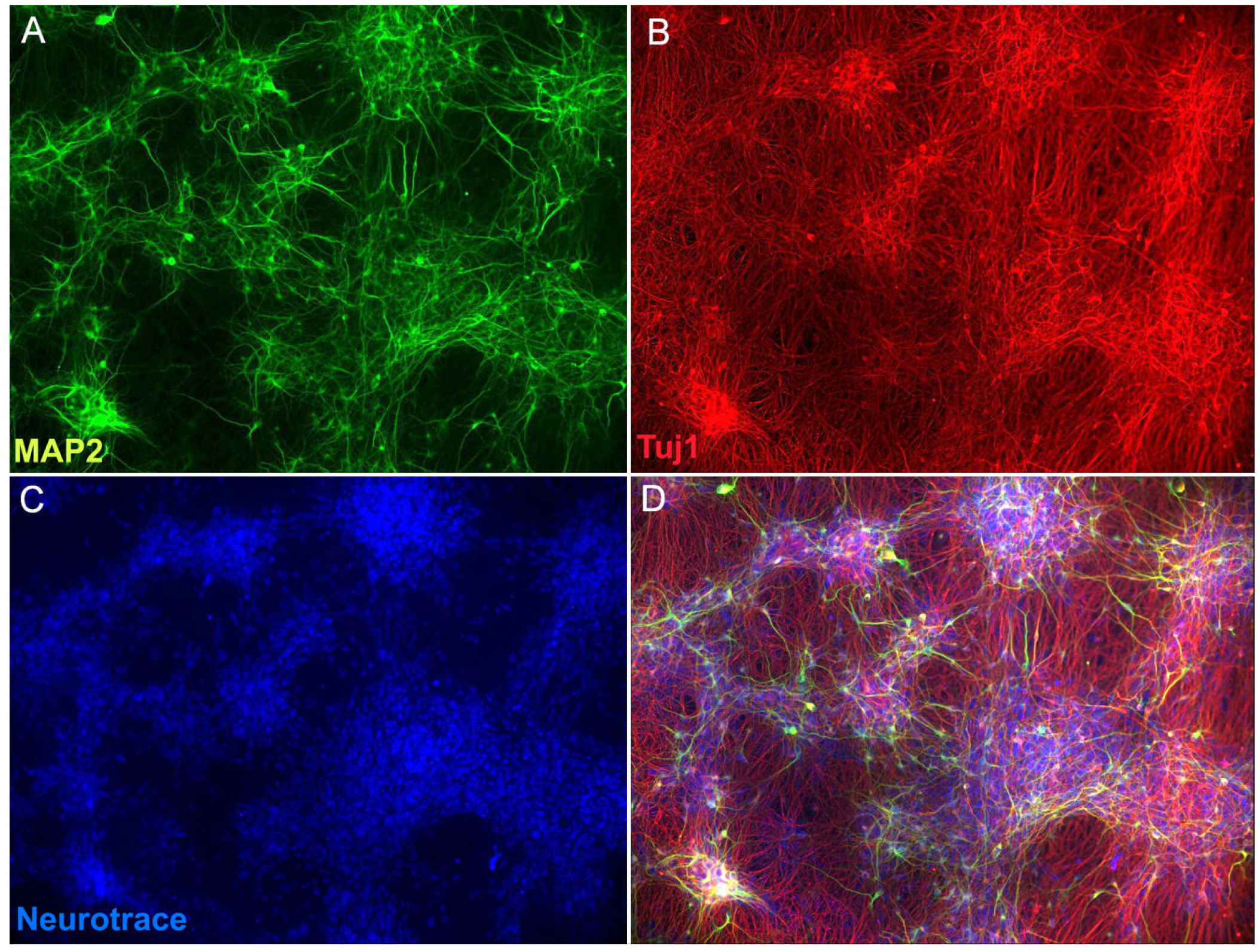
iPSC-neurons (iPSC-neurons) are express neuronal markers. Immunostaining for: (A) MAP2 (green), (B) Tuj1 (red), and (C) Neurotrace (blue), (D) merge.

**Figure 2.**
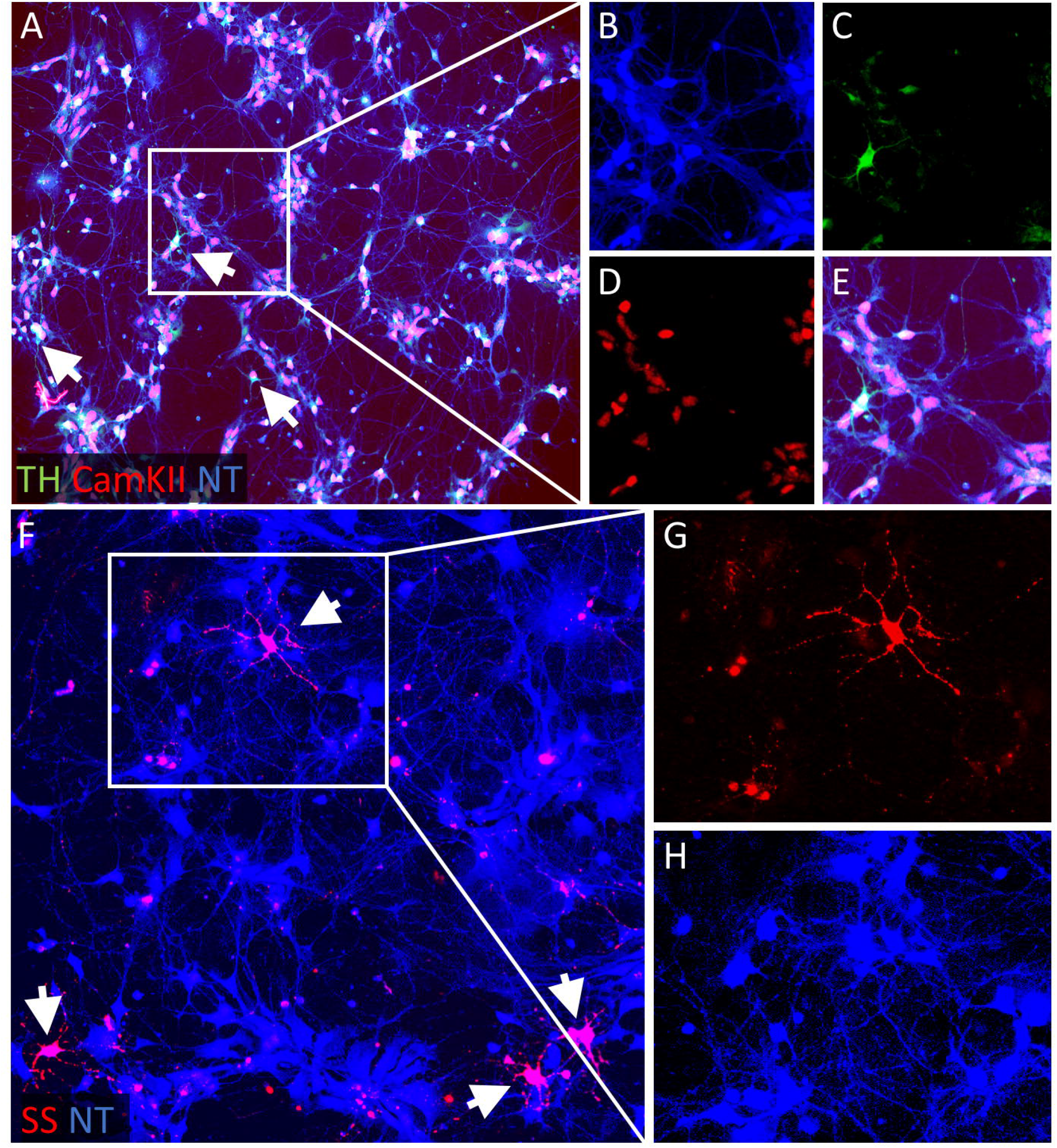
A heterogeneous populations of iPSC-neurons. A-E CAMKII (red) was used to detect glutamatergic and tryrosine hydroxylase (TH) (green, arrows) was used to detect dopaminergic neurons. B-E are a magnification of the cutout window in A. Neurotrace (NT) (blue) was used as a neuronal counterstain. (F-H) Somatostatin (SS) staining revealed a subgroup of GABAergic neurons (arrows). G,H are a magnification of the cutout window in F. 80%, 0.58%, and 4.8% of Neurotrace+ cells (blue) were colabeled with each of the markers, respectively.

To assess whether human iPSC-neurons expressed receptors of interest, we measured mRNA levels of our receptors of interest over the course of neuronal differentiation. We found that each of the receptors assessed exhibited their own temporal profile of expression. Because glutamate is the major neurotransmitter in the brain, we anticipated abundant glutamate receptor expression in our iPSC-neurons. Indeed, we found a significant increase in mRNA of glutamate receptors, including kainate (GRIK1), and NMDA receptor subunit 1 (GRIN1) following 79 and 115 days of differentiation (Fig 3). Remarkably, the onset of GRIK1 and GRIN1 expression in human iPSC-neurons was consistent with the Allen BrainAtlas Developmental Transcriptome, which demonstrates increased expression of these receptors as early as 8 weeks post conception in human fetal tissue (Miller et al., 2014; © 2016 Allen Institute for Brain Science). In contrast, we found no significant changes in NMDA receptor subunit 2A (GRIN2A) expression at the time periods assessed, up to 115 days, which is equivalent to approximately 16 weeks post conception. Importantly, GRIN2A expression does not increase in humans (Miller et al., 2014; © 2016 Allen Institute for Brain Science) or rodents (Liu et al., 2004) until birth. Thus, in line with previous reports (Shi et al., 2012; Brennand et al., 2015), neuronal differentiation of human iPSCs by Dual-SMAD inhibition recapitulates aspects of human brain development.

**Figure 3.**
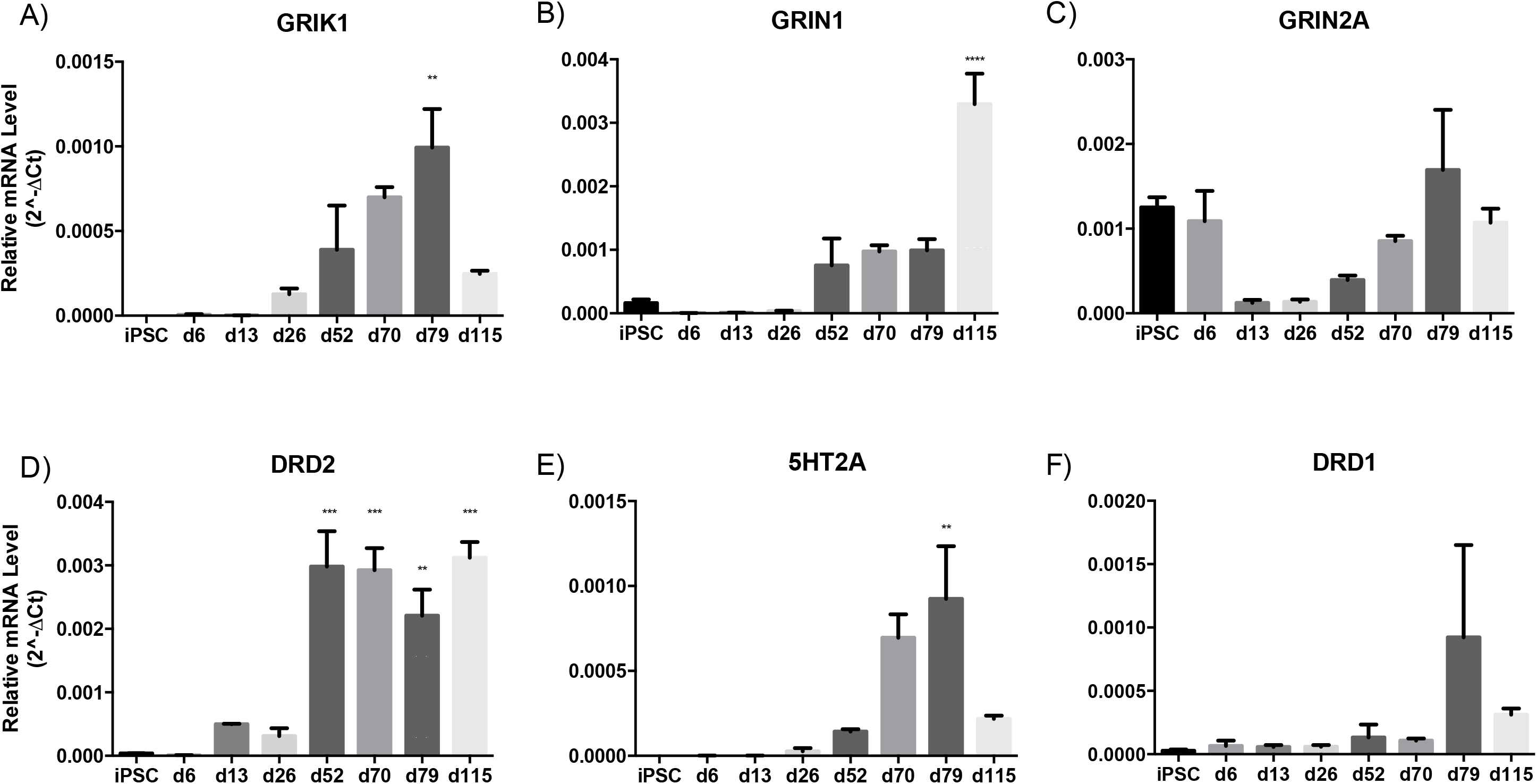
Receptor expression over the course of neuronal differentiation of human iPSCs. mRNA levels are represented as 2^^-ΔCt^ normalized to housekeeping genes actin and GAPDH. All values are expressed as the mean ± SEM, n=3-6. Statistical analysis was performed on Prism 6 (GraphPad) using one way AN OVA followed by Dunnett’s multiple comparison post hoc test compared to iPSC group. p<0.05*, p<0.01**, p<0.001***, p<0.0001****

Prior studies using human iPSC-neurons have demonstrated gene expression for glutamate and GABA receptors (Dage et al., 2014). Here, we report that iPSC-neurons express modulatory amine receptors, which serve as targets for antipsychotic medications. Atypical antipsychotic drugs bind to both D2R and 5HT2AR amongst numerous other receptors. We found that differentiating neurons increase expression of D2R by day 52 of differentiation (Fig 3) and that the increased expression was maintained through day 115. Human iPSC-neurons also expressed higher levels of 5HT2AR on day 79 of differentiation (Fig 3) that decreased by day 115. We did not find significant expression of D1R in our iPSC-neurons (Fig 3).

Clozapine is the oldest and most efficacious atypical antipsychotic and binds to the D2R, D1R, and 5HT2AR amongst other receptors (Meltzer, 2012). To understand how an antipsychotic such as clozapine might modulate neuronal activity, neurons were focally stimulated with glutamate, dopamine, and 5-HT2A agonist DOI, before and after incubation with clozapine or vehicle. Calcium measurements in iPSC-neurons provide detailed functional information about neuronal phenotypes following differentiation (Glaser et al., 2016). Therefore, cells were preloaded with a fluorescent calcium indicator, and changes in calcium levels were recorded to monitor neuronal activity in response to the pharmacological challenges. Neurons were identified by morphology in brightfield, and calcium responses of individual neurons were analyzed and grouped into “subtypes” depending on their individual response profile. A 10% increase over baseline calcium fluorescence levels has been correlated to single action potential spiking (Kerr et al., 2005; Sasaki et al., 2008), and corresponded to a consistent visible calcium flux following drug application in our system. Therefore, using this cutoff as a measure of neuronal response, we compared response patterns of cells exposed to vehicle solution (Group A) to those exposed to clozapine (Group B).

Based on our criteria, we found four major neuronal response patterns. Approximately half of the neurons exposed only to vehicle or following clozapine slow bath solution only responded to glutamate (Groups A and B, Fig 4A). These cells may represent a subpopulation of neurons with functional glutamate receptors, but not dopamine or 5HT2A receptors, and suggests that neurons with this response profile do not change calcium signaling in response to clozapine incubation. Interestingly, some neurons responded to an initial, but not second stimulation with glutamate (Fig 4B). Remarkably, incubation with clozapine decreased the number of neurons that failed to meet criteria for mounting calcium transients upon re-exposure to glutamate (Fig 4B) suggesting that clozapine increased the likelihood of calcium transients in response to glutamate exposure. Conversely, some neurons mounted calcium transients not on initial, but only on a repeat exposure to glutamate (Fig 4C). This population was also more prevalent after clozapine treatment (Fig 4C) suggesting that clozapine treatment can increase the likelihood of glutamate-induced calcium transients even in neurons that do not normally respond to glutamate. Together these findings suggest that clozapine may potentiate glutamate responses in neurons that are normally sub-threshold to mount a calcium response to glutamate.

**Figure 4.**
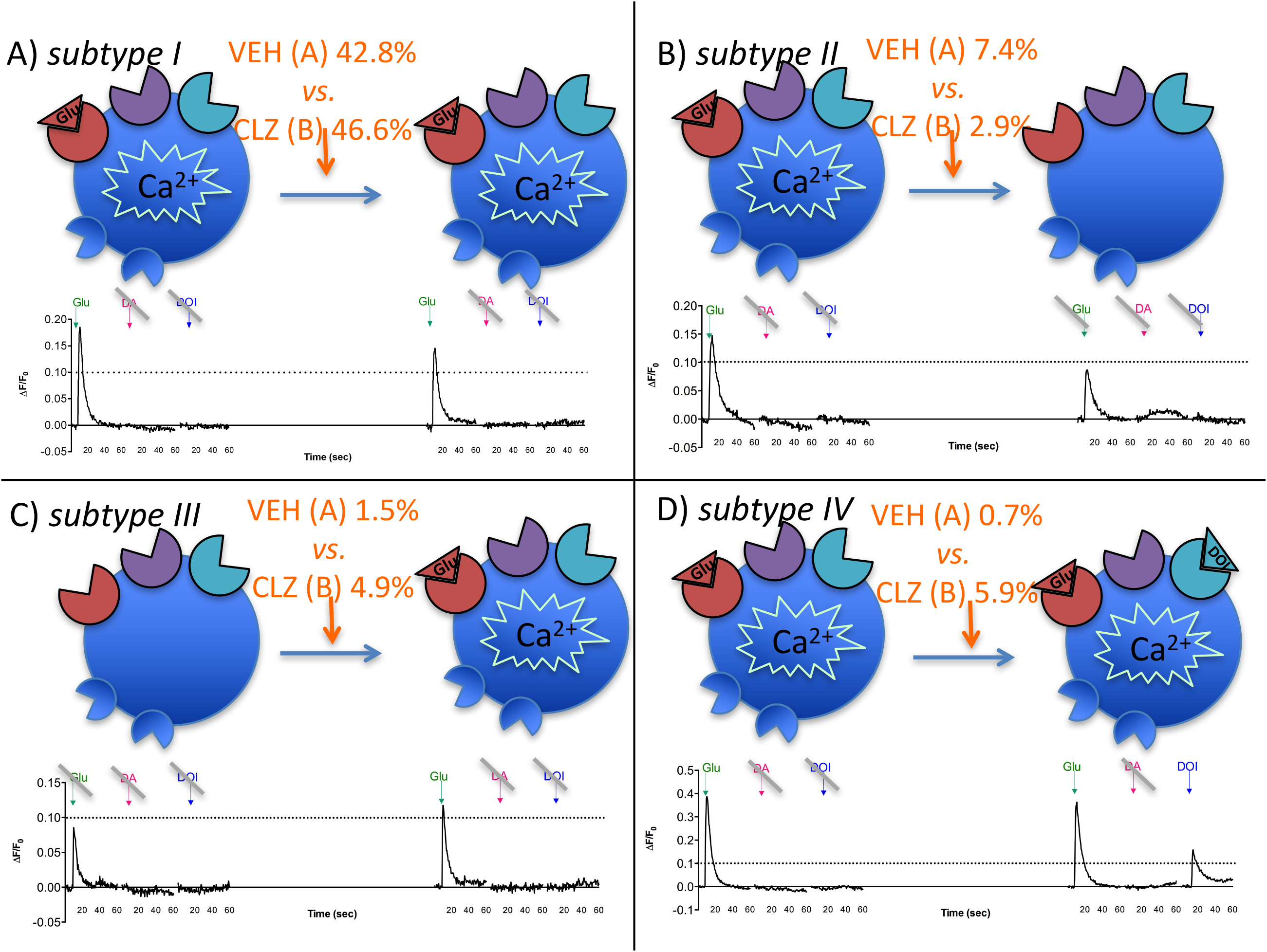
Calcium response profiles of individual neurons. Calcium levels were monitored following evoked focal stimulation with glutamate (Glu), dopamine (DA), and 5HT2A agonist (DOI) before and after exposure to a slow bath perfusion of vehicle (Group A) or clozapine (Group B) solution. Neurons were analyzed and a calcium increase 10% above baseline (indicated by dotted line) was defined as a response. Distinct calcium response profiles emerged: (A) Subtype I neurons responded to an initial and second stimulation with Glu; (B) Subtype II neurons responded to an initial Glu stimulation; (C) Subtype III responded to a second Glu stimulation; (D) Subtype III neurons responded to Glu and a second stimulation with DOI. Representative calcium traces are shown below each subtype cartoon. (Group A vehicle slow bath: n=271, Group B clozapine slow bath: n=410).

Very few cells mounted calcium transients in response to dopamine or serotonin receptor targeting. One such notable subtype of neurons responded initially only to glutamate, but then to glutamate and the 5HT2A agonist DOI upon re-exposure (Fig 4D). Remarkably, iPSC-neurons exhibiting this last pharmacological response profile were dramatically increased by clozapine treatment (Fig 4D). This finding suggests that clozapine may enhance neuronal responses via 5HT2AR signaling that occurs following glutamate stimulation. It was rare to find neurons that responded to DOI exclusively, with dopamine, or other combinations with glutamate (Suppl Table 1). Moreover, there were also many neurons that did not elicit responses to any of the tested ligands (~44% and 33% in Group A and B, respectively). All other unique response profiles constituted less than 2% of the cellular population in both Groups A & B (Suppl Table 1). Together our findings demonstrate heterogeneity of individual neuronal calcium responses to neurotransmitters and suggest that clozapine modulates calcium responses to glutamate and serotonin signaling. These results also suggest that single neuron pharmacological profiling is an important consideration when analyzing cellular antipsychotic drug response.

## DISCUSSION

The current studies support the use of human iPSC-neurons to investigate antipsychotic drug pharmacology. Using the general Dual-SMAD inhibition protocol to differentiate iPSCs, we found evidence to support a mostly glutamatergic population, and much lower levels of GABAergic and dopaminergic neuronal expression (Fig 2). This is consistent with previous reports suggesting that using this method mimics forebrain development (Brennand et al., 2011; Shcheglovitov et al., 2013).

To understand whether iPSC-neurons present a viable tool for studying antipsychotic pharmacology, we first probed expression of relevant receptors. We found that D2R and 5HT2AR were not expressed until more than a month into the differentiation process in our iPSC-neurons (Fig 3). This developmental trajectory was expected for D2R, but surprising for 5HT2AR, which are not normally expressed until approximately 4 months in the prefrontal, temporal, and primary visual cortex(© 2016 Allen Institute for Brain Science). Overall, our results generally support previous studies using human iPSC-neurons that show gene expression profiles closest to first trimester fetuses, from 8 to 16 weeks post conception (Brennand et al., 2015), but extend the findings to expression of functional receptors. Another study has demonstrated that iPSC-neurons follow the time line of human corticogenesis, spanning approximately 70 days in comparison to 6 days using mouse iPSCs (Shi et al., 2012). Thus, human iPSC-neurons have validity in reflecting human neuronal development especially for neurotransmitter receptors. While this quality is a strength of the system, it also poses a limitation because it is unclear whether receptor expression at these early stages is relevant to later adult psychopharmacology. Future studies using iPSC-neurons from patients can be used to identify whether early receptor expression profiles differentiate patients with different drug responses.

Following evidence for receptor expression, we sought to determine whether these receptors were functional and whether clozapine could modulate their responses to glutamate, dopamine, and the 5HT2AR agonist (DOI). Single cell response profiling revealed changes in the proportion of neurons in several subtype response groups when exposed to clozapine compared to vehicle slow bath solution (Fig 3). We found that clozapine did not alter glutamatergic responses in most cells. Remarkably, in subpopulations of cells clozapine treatment modulated glutamate responses, potentiating them in some neurons, while attenuating them in others (Fig 3, subtypes II and III). Our results therefore suggest that clozapine can bi-directionally modulate glutamate transmission in neuronal subpopulations. The biology distinguishing these subpopulations remains to be determined. However, paradoxical responses in a seemingly homogeneous cellular population highlights the importance of single cell analysis in pharmacological studies.

Remarkably, clozapine induced an otherwise rare DOI response in some iPSC-neurons. These cells (Group IV) were distinct from other groups in that clozapine did not appear to modulate response to DOI as it did for glutamate in Groups II and III. Instead, Group IV neurons exhibited an otherwise absent calcium response to DOI indicating a more qualitative difference. Evidence suggests that clozapine induces increased expression of 5HT2AR in rodent cortical pyramidal neurons (Willins et al., 1998) highlighting one possible mechanism, by which DOI responses of some iPSC-neurons were altered. Regardless of the mechanism for this specific responsiveness change induced by clozapine, our results suggest that mixed iPSC-neuron preparations provide a viable platform for mechanistic pharmacologic studies. This platform is unique in that it provides a clear path to inquiry of how serotonin antagonism may contribute to clozapine’s and other atypical antipsychotics’ clinical efficacy. Such a path would include iPSC-neurons derived from patients with differential antipsychotic drug response to identify cellular phenotypes for personalized medicine.

Here we report the feasibility of generating neurons from iPSCs for the use in psychopharmacology studies. These are the first studies to directly assess the presence and function of D2R and 5HT2AR in human iPSC-neurons, and modulation of these receptor activities via clozapine. Future studies should include alternative differentiation protocols that enrich for dopaminergic or GABAergic neurons to understand how antipsychotics might differentially affect large populations of these neuronal subtypes. In addition, the use of lentiviruses with fluorescent probes to mark neuronal subtypes combined with pharmacological profiling may allow for better characterization of current antipsychotics and targeting of future therapies. Importantly, identifying a small population of cells exhibiting a categorically different response to an efficacious medication, raises an exciting possibility that yet uncharacterized cellular subpopulations may serve as distinct psychopharmacological targets.

## METHODS

### Neuronal differentiation

Control human iPSC cell line CDI iPSC 8325 was kindly provided by the NIH Center for Regenerative Medicine. This line was derived from human peripheral blood mononuclear cells and reprogrammed using episomal methods. iPSCs were differentiated using Dual-SMAD inhibition based on published methods (Tomishima, 2008) with minor modifications. On day 10-13 differentiation, cells were passaged with Accutase onto polyornithine/laminin/fibronectin coated plates. Neuronal differentiation media consisted of neurobasal media supplemented with N2, B-27, and Glutamax, and fresh BDNF (20ng/ml) and ascorbic acid (200uM) was added to the media immediately before use. Cells for calcium imaging were plated onto coated 15mm coverglasses in 24-well plates, and allowed to recover for 2 weeks before imaging.

### Immunocytochemistry

Half of cell media was removed, and prefixed at a final concentration of 2% PFA for 10 min, followed by 4% PFA fixation for 10 min. Cells were washed 3X with PBS for 5 min each. Cells were blocked (0.1% Triton, 10% NDS) at RT for 1 hr, and then incubated with primary antibodies: MAP2 (Sigma M9942), TH (Pelfreez P60101-0), CAMKII (Millipore 05-532) diluted at 1:500 in blocking solution (0.1% Triton, 1% NDS) overnight, shaking at 4°C. Cells were then washed 3X with PBS for 5 min each at RT on a shaker. Secondary antibodies were diluted in blocking solution (1:500) and added for 1 hr at RT, covered and shaking. Cells were washed 3X with PBS, 10 min each., incubated with Neurotrace (Life Technologies N-21479) at 1:400 for 10 min and washed 3X. Glass coverslips were carefully removed, rinsed once with H_2_O, and mounted onto slides.

### Lentiviral Transduction

Human synapsin-RFP lentivirus was generously provided by Dr. Derek Dykxhoorn at University of Miami. Cells were transduced with 4.85 x 10^8^ IU/ml with an MOI of 5. RFP positive neuronal cell bodies and processes were visible within 3 days and imaged using an Olympus epifluorescent microscope.

### Quantitative RT-PCR

RNA was harvested using Trizol Reagent (Invitrogen) according to manufacturer’s protocol. cDNA was synthesized using SuperScript^®^ III Reverse Transcriptase (Invitrogen) from 3ug RNA. Primers were designed using Primer 3 and human mRNA sequences from NCBI as follows: GRIK1 F: cctcggtggacaacaaagat R: gctggcggattttgatttta; GRIN1 F: ccaagcccttcaagtaccag R: ctcctcctcgctgttcacct; GRIN2A F: ggttgctcttctccatcagc R: gcagctcttttgggtgagtc; DRD2 F: gatctttgagatccagaccat R: atgttgcagtcacagtgtatgt; 5HT2A F: tcatcatggcagtgtcccta R: gagcacgtccaggtaaatcc; DRD1 F: gaattgccagaccaccacag R: ccacccaaaccacacaaaca. qPCR was performed using *Power* SYBR^®^ Green Master Mix and StepOnePlus™ Real-Time PCR System (Applied Biosystems). mRNA levels are represented as 2^^-ΔΔCt^ relative to the iPSC group and normalized to housekeeping genes actin and GAPDH. All values are expressed as the mean ± SEM, n=3-6. Statistical analysis was performed on Prism 6 (GraphPad) using one way ANOVA followed by Dunnett’s multiple comparison post hoc test to compare each day of differentiation to iPSC.

### Calcium Imaging

Cells were seeded onto glass coverslips coated with polyornithine/laminin/fibronectin 17 days prior to calcium imaging. Fluo-4 was reconstituted with DMSO to a 5mM stock, and diluted 1:1 right before use with Pluronic F-127. At 68 days of differentiation, cells were washed twice with HEPES recording buffer (145mM NaCl, 5mM KCl, 10mM HEPES, 2mM CaCl_2_, 2mM MgCl_2_, 10mM glucose), and loaded with a final concentration of 2uM Fluo-4 for 45 min at RT. Cells were rinsed twice with HEPES buffer and incubated for 30 min to equilibrate and for de-esterification. Coverslips with cells were loaded into a Warner perfusion chamber, and calcium imaging was performed using a Nikon Eclipse TE300 inverted microscope with pco.EDGE CMOS camera (pco, Germany) and 20X objective. Brightfield DIC image was first taken to identify a field of view. Calcium transients were recorded in the FITC filter, and images were acquired at 2 Hz. A slow bath perfusion of HEPES buffer flowed over the coverslip during the entire recording. For ligand evoked calcium recordings, a microtube (~0.3mm diameter) was positioned directly adjacent to the field of view and released select ligands: glutamate, dopamine, or 5-HT2A agonist DOI hydrochloride (100μM each) 6 sec after the beginning of the recording. Each ligand was released for 2 sec, followed by buffer solution for the remainder of the one minute recording. CamWare V3.10 and Micromanager software programs were used to release solutions from the microtube and record evoked calcium transients. A buffer period of at least 3 minutes of vehicle slow bath perfusion was performed between each recording. Maintaining the same field of view of cells, either 1 μM clozapine (Sigma) or vehicle solution was added to the slow bath perfusion for 10 minutes, followed by stimulation with glutamate, dopamine or DOI. Neurons were analyzed before and after perfusion with clozapine or vehicle solution.

### Calcium Imaging Analysis

Following acquisition, images were analyzed using Fiji software. Neurons were identified by morphology as a cell body with one or more processes and regions of interest (ROIs) were manually drawn. The mean grey values of ROIs during the recording were obtained, and neurons with a 10% increase in ΔF = (F-F0)/F0 were defined as cells that responded to evoked stimulation based on previous studies(Kerr et al., 2005; Sasaki et al., 2008) and the ability to detect calcium fluxes using this cutoff in our system. Responses of individual cells after evoked stimulation to glutamate, dopamine, and DOI were examined before and after clozapine (Group A) or vehicle (Group B) slow bath perfusion. We analyzed 271 and 410 neurons for each group, respectively. Each neuron was classified into a response profile based on response to the select ligands. Response profiles shared by <2% of the cells were considered rare events and not included as a major subtype.

## Acknowledgments

We would like to thank Andrew Sproul and Aiqun Li at the New York Stem Cell Foundation, and Damian Williams, Barabara Corneo, and Alejandro Garcia Diaz at the Columbia Stem Cell Facility for their technical support and help and Derek Dykxhoorn at the University of Miami for sharing lentiviruses. ESK is supported by 5T32MH016434 and the Dr. Joseph & Lillian Pisetsky Award. This work was supported by NIMH R01MH091844 (to AD), R01MH105675 (to EDL) and the Irving Scholar award from Columbia University (to AD).

